# Mechanistic underpinning of nonuniform collective motions in swarming bacteria

**DOI:** 10.1101/2021.02.08.430357

**Authors:** Palash Bera, Abdul Wasim, Jagannath Mondal, Pushpita Ghosh

**Author notes:** Electronic address (Pushpita Ghosh).

## Abstract

Self-propelled bacteria can exhibit a large variety of non-equilibrium self-organized phenomena. Swarming is one such fascinating dynamical scenario where a number of motile individuals grouped into clusters and move in synchronized flows and vortices. While precedent investigations in rod-like particles confirm that increased aspect-ratio promotes alignment and order, recent experimental studies in bacteria *Bacillus subtilis* show a non-monotonic dependence of cell-aspect ratio on their swarming motion. Here, by computer simulations of an agent-based model of self-propelled, mechanically interacting, rod-shaped bacteria in overdamped condition, we explore the collective dynamics of bacterial swarm subjected to a variation of cell-aspect ratio. When modeled with an identical self-propulsion speed across a diverse range of cell aspect ratio, simulations demonstrate that both shorter and longer bacteria exhibit slow dynamics whereas the fastest speed is obtained at an intermediate aspect ratio. Our investigation highlights that the origin of this observed non-monotonic trend of bacterial speed and vorticity with cell-aspect ratio is rooted in the cell-size dependence of motility force. The swarming features remain robust for a wide range of surface density of the cells, whereas asymmetry in friction attributes a distinct effect. Our analysis identifies that at an intermediate aspect ratio, an optimum cell size and motility force promote alignment, which reinforces the mechanical interactions among neighboring cells leading to the overall fastest motion. Mechanistic underpinning of the collective motions reveals that it is a joint venture of the short-range repulsive and the size-dependent motility forces, which determines the characteristics of swarming.

## I. INTRODUCTION

Collective motions and self-organized order formation at the expense of internal or external sources of energy are commonly observed in many living systems across a wide length scales. Starting from the coherent motion of cells in tissue flow, motion in bacteria, fish, and birds, this also holds for collective migration of animals [1–4]. In micro-organisms, self-propulsion is a form of active motion that is responsible for the emergence of various orders in cell-collectives such as motility-induced phase-separation [5], chemotaxis [6–8], pattern formation [9–14], swarming and turbulent motions [15–20]. Bacterial swarming is an efficient mode of collective migration of a group of densely-packed, rod-shaped flagellated bacteria on the surface. During swarming, collectively moving dynamic clusters of cells exhibit coherent flows, swirls, vortices that can persist for several seconds [21, 22]. In highly dense bacteria colonies, mechanical interaction becomes relevant and the tendency for neighboring particles to align is strongly determined by their mutual interactions [23, 24]. The underlying dynamics of the swarming phenomenon stems from the self-propelled motion of individual rod-shaped cells along with the short-range steric forces [18, 25, 26]. However in some instances, long-range hydrodynamic interactions might be relevant [27, 28] to determine the self-organized motion.

Over the last few years, there has been extensive research and progress in understanding the collective motions and the underlying dynamics in connection with the biological manifestation of cell-elongation, increased flagella density during swarming motion [15, 29]. A number of prior studies have focused on mainly understanding the swarming dynamics with respect to the properties like cell-density or packing fraction and cell-aspect ratio, constant and density-dependent self-propulsion forces, a mixture of motile and non-motile cells [13, 17, 30–33]. For a statistical physics view on bacterial swarming dynamics we refer [34]. The majority of the previous studies have mainly focused on velocity and vorticity distributions, relation with time and length scales, cluster-size distributions, spatial and temporal correlations during swarming dynamics [16, 18, 35–39]. In this regard, investigation on bacteria *Bacillus subtilis* by Ilkanaiv and co-workers [32], have reported that swarming statistics depends on the cell-aspect ratio in a critical and significant way not described by theory. Both shorter and longer cells show low mobility and non-Gaussian statistics whereas intermediate or wild-type shows the fastest mobility and Gaussian behavior. Recently, they also have shed light on the collective dynamics of a monolayer of bacterial cells with respect to the variation of surface density and cell aspect ratio [33]. While precedent investigations in asymmetric rod-like particles confirms aspect ratio dependence in their collective dynamics, underlying mechanistic insights are still lacking. As cell-size has a profound impact on cellular-organization, a variation of cell aspect ratio invites a fundamental question: How does the cell-aspect ratio contribute to microbial systems? Does cell-size play a role in beneficial movement or efficient collective motion in a group?

In a swarming bacteria colony, the feature which is of prime importance is the self-propulsive motion of individual cells. The motion of an individual bacteria in a dilute suspension where neighboring interactions are absent is mainly stems from its self-propulsive force. However, it has been indicated in [32, 33] that self-propulsion speed at individual level is nearly the same for all the different mutants irrespective of their cell aspect ratios. This feature has not been considered in some of the prior studies [17, 40] which dealt only with constant motility forces. Our aim here is to emphasize on this key feature of motile cells and establish the connection between aspect ratio and motility. Using an individual/agent-based model of rod-shaped bacteria following an overdamped dynamics in a monolayer of swarm, we have shown that the dependence of self-propulsion force on cell-size might be a crucial factor that spontaneously gives rise to a non-monotonic dependence of mean-speed and vorticity on aspect ratio of bacteria in swarming colonies. We carry out several control simulations and analysis of the total translational and rotational forces on each cell to distinguish between the contributions of the two main forces on the cellular motions. Our results illuminate that an interplay of the dependence of cell-size on motility and the steric repulsive interactions among the neighboring bacterial cells leads to the fastest motion in the case of cells having an intermediate cell aspect ratio. Both shorter and longer cells exhibit slow dynamics as observed by their mean speed and velocity autocorrelation functions. Moreover, we show that asymmetry in friction has distinct effect in cell motion which might enhance the mean speed of the colony when rotating motion is easier than translational. Our study is relevant to a class of motile asymmetric particles which differs in their cell-size but posses nearly the same self-propulsion speed, follows simple over-damped dynamics where short range cell-cell mechanical interactions are dominant over hydrodynamics and is able to provide sufficient insights on their nonuniform collective statistical dynamical behavior.

## II. SIMULATION MODEL AND METHOD

We consider an individual-based model of self-propelled bacterial cells in two-dimensions. Each bacterial cell is modeled as motile spherocylindrical particle of constant diameter or width (*d*_0_ = 1*μ*m) and length *L* = *l* + *d*_0_, where *l* corresponds to the cylindrical length of an individual cell and depicted in Figure 1(a).

**FIG. 1:**
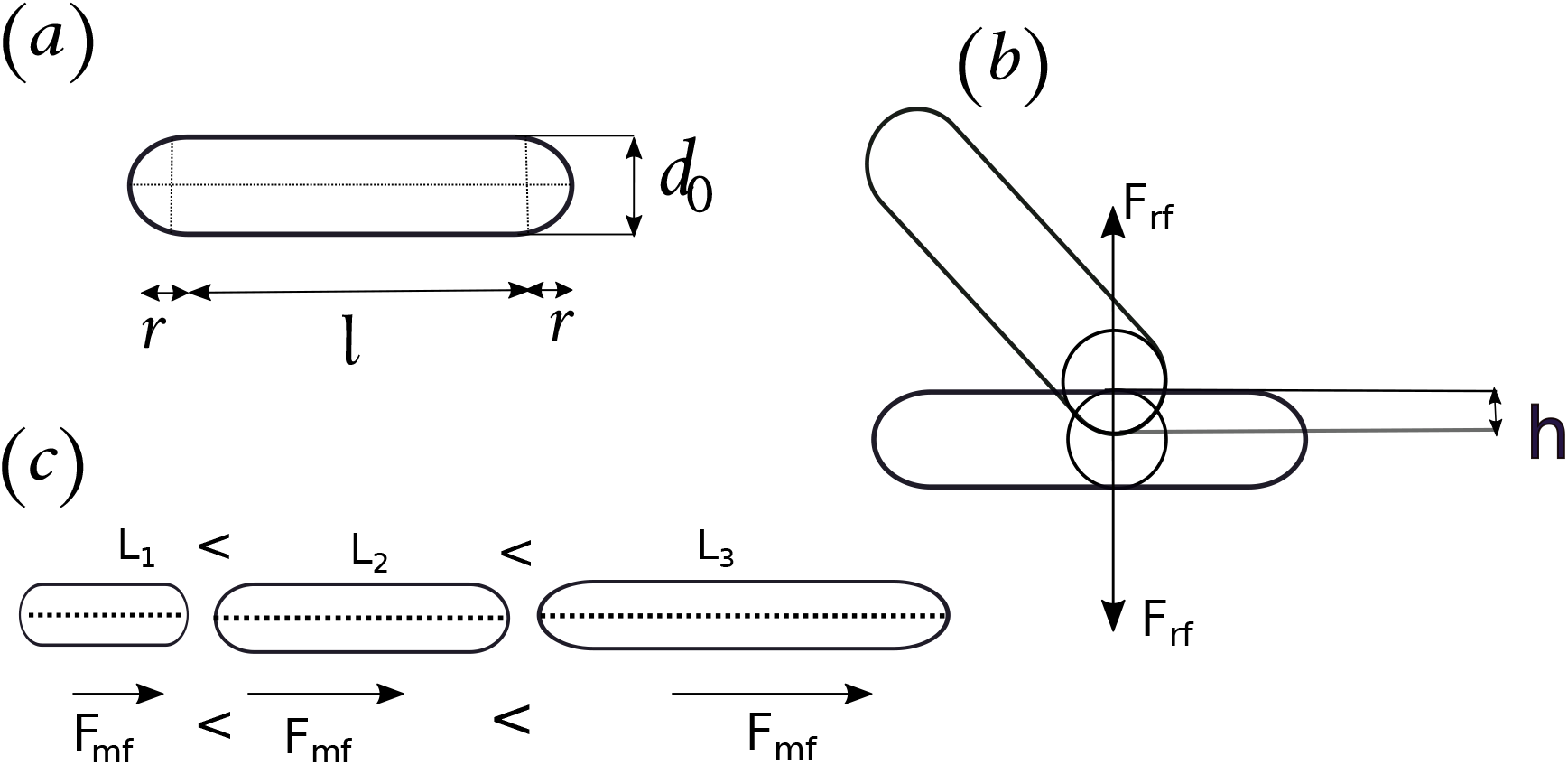
(a) Shape of bacteria as spherocylinders with a cylindrical length *l* and radius of the end caps *r,* (b) Cell-cell repulsive mechanical interaction between two bacteria, (c) The motility force, *F_mf_* increases with increasing aspect ratio and is directed about the major axis of the cell.

The equation of motions for the bacterial cells follow an over-damped dynamics. The latter assumption is justified for a densely packed colony where viscosity dominates over inertia. We assume that the particles are submerged in a viscous medium such that the linear velocity and the angular velocity are proportional to the force and torque acted upon a rod-shaped cell respectively as described by the following equation of motion.

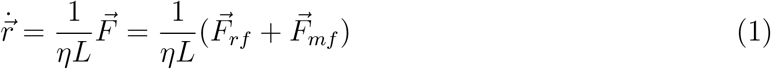

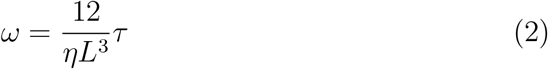

where 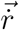 and *ω* are linear and angular velocities respectively and *η* represents the friction coefficient. Even though the rod shape of the particles requires different friction coefficients that correspond to resistance exerted by the surrounding medium when particles either rotate or move along their short or long axes but for simplicity, we assume the same friction coefficient (*η*) in all directions at first and as we will divulge into the result section, we will explore the impact of having different frictions on the results. At each time step of the simulation, we systematically corrected the center of mass velocity of the simulation box, thereby ensuring is no external force acting on the system.

As shown in equation 1, each bacterial cell is equipped with a motility force resulting in a self-propelled velocity acting along its long axis. In addition, we consider a shortrange repulsive mechanical interaction among the neighboring cells, neglecting the long-range hydrodynamic interactions. Accordingly, the model describes a net force acting on each self-propelled bacterial cell as the summation of a repulsive component of the force (*F_rf_*) and a component attributing to the motility-force (*F_mf_*).

We assume that cells interact mechanically through repulsive interaction which is in accordance with the Hertzian theory of elastic contact. We adopt the same functional form of *F_rf_* as employed in previous studies [23, 24, 41, 42] and is given by: 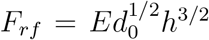, where *E* represents the elastic modulus of the cells and *h* corresponds to the overlap between two interacting cells. Figure 1(b) schematically demonstrates the repulsive mechanical interaction between two bacteria where *h* = *d*_0_ — *r*_0_ and *r*_0_ is the closest distance of approach between the two rod-shaped cells.

Apart from the cell-cell repulsive mechanical interaction, the essential feature of an individual bacterium is the presence of self-propulsive or motility force (*F_mf_*), which is one of the key points in our current study. The experimental studies in references [32, 33], have suggested that the swimming speed of bacteria in very sparse liquid suspensions, where bacteria hardly interact with each other and with no interaction with a surface, is nearly the same for all the mutant strains having different aspect ratios. On the other hand, as per Eq.(1), since under over-damped condition, force is proportional to the velocity, it suggests that to have the same self-propulsion speed as smaller body, larger motility force is needed to move the longer body. Based on this feature, it turns out that the motility force might depend on cell-length in some characteristic manner. In this regard, in our model, we propose that the motility force is a linear function of cell-length i.e. *(F_mf_* ℝ *L*, so that each individual bacteria has the same self-propulsion velocity irrespective of their cell aspect ratio under an over-damped condition. In other words, the flagella motor speed linearly increases with the increase of the cell-aspect ratio. A schematic representation of motility force *F_mf_* directed about the major axis of the cells are shown in Figure 1(c). However, the actual functional form linking the motility force with the cell size is not precisely known and the underlying biological origin might be more complex. By this modulation of motility force, under the aforementioned conditions, we simulated the spatiotemporal dynamics of bacterial colonies at a range of cell aspect ratios. The generic functional forms of the motility force expression employed in the current work can be expressed as *F_mf_* = *f_mot_ L* where *f_mot_* refers to the proportionality constant.

Table I enumerates the values or range of parameters employed in the current investigation. There are four key parameters that control the swarming dynamics: (i) motility force, (ii) the packing fraction *ρ*, i.e the area occupied by the particles divided by the total area (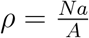, where *N* is the number of the particles in the simulation box, *a* corresponds to the area of a single particle and *A* is the total area of the box), (iii) the aspect ratio of the bacteria (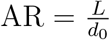, where *L* is the length and *d*_0_ is the width/diameter of the spherocylindrical cells), and (iv) the value elastic modulus (E).

**TABLE I:**
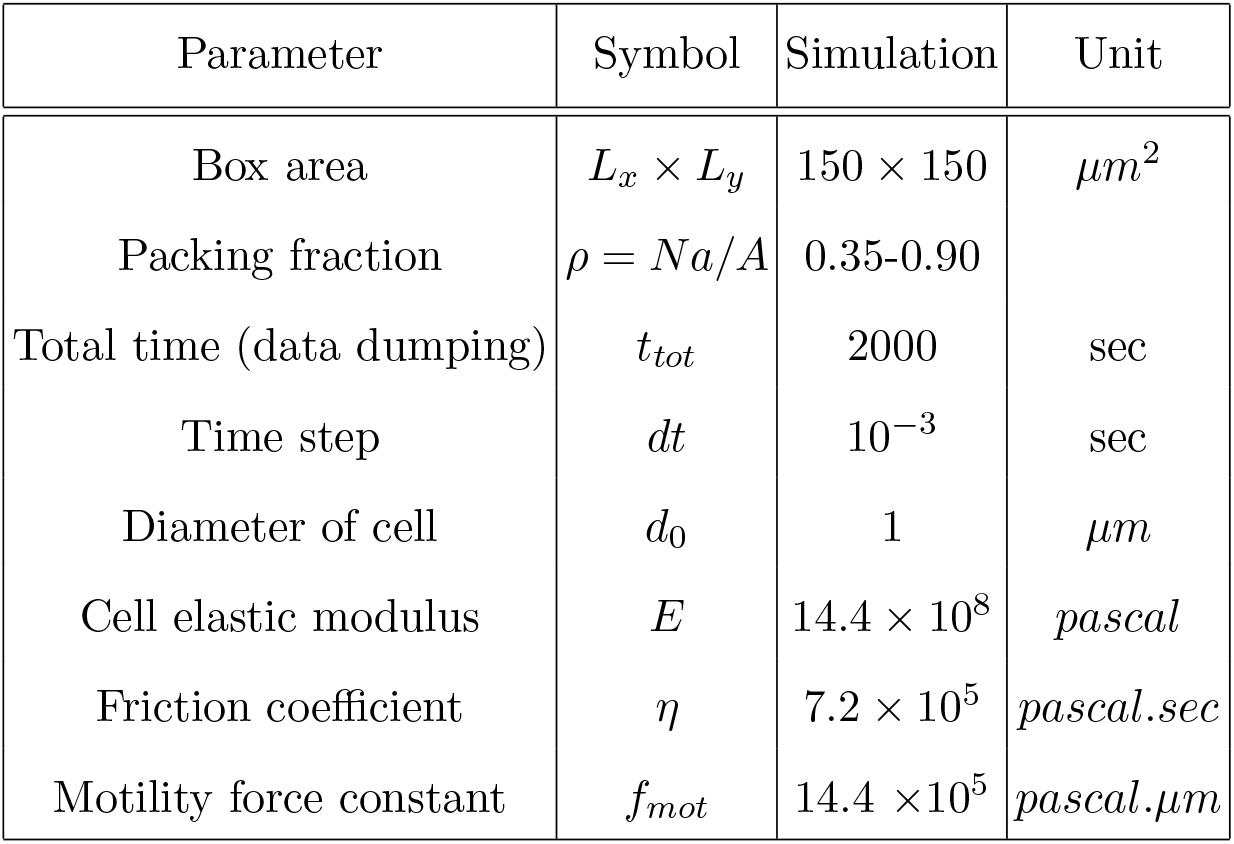
Parameters and constants used in our agent-based model

| Parameter | Symbol | Simulation | Unit |
| --- | --- | --- | --- |
| Box area | $L_x \times L_y$ | $150 \times 150$ | $\mu m^2$ |
| Packing fraction | $\rho = Na/A$ | 0.35-0.90 | |
| Total time (data dumping) | $t_{tot}$ | 2000 | sec |
| Time step | $dt$ | $10^{-3}$ | sec |
| Diameter of cell | $d_0$ | 1 | $\mu m$ |
| Cell elastic modulus | $E$ | $14.4 \times 10^8$ | *pascal* |
| Friction coefficient | $\eta$ | $7.2 \times 10^5$ | *pascal.sec* |
| Motility force constant | $f_{mot}$ | $14.4 \times 10^5$ | *pascal.$\mu m$* |

To create a spontaneously formed bacterial colony, we initiated each of the simulations via seeding with a few numbers of cell particles (~ 4) randomly placed inside the simulation box. Subsequently, each bacterial cell grows at a constant uniform rate (*γ*=0.2/sec) and subsequently splits into a pair of daughter cells when the original cell reaches its maximum threshold length of division. When the colony, thus formed, reaches the desired packing fraction (*ρ*), the cell-division stops. However, at this time, if there are cells remaining which are yet to reach their maximum length, these will still grow and finally stop once they meet the criteria of the identical aspect ratio for each individual in the colony. This condition mimics the experimental setup of typical growing colony of a large number of densely packed cells before they reach the steady-state condition. This approach ensures that the modeled bacterial cells arrange naturally in a dense-pack arrangement, just like in a growing bacterial colony.

## III. RESULTS AND DISCUSSION

The key objective of the present study is to explore the spatiotemporal dynamics of rodshaped bacteria in dense colonies those differ by their cell aspect ratios which in essence is motivated by the experimental studies [32, 33] of swarming dynamics observed in *B. subtilis*. Cell aspect ratio deserves a special attention as an important parameter, because some of the bacterial species change their cell aspect ratio before starting to swarm, suggesting that the aspect ratio is key to swarming dynamics [15, 29]. Our investigation proposes that bacterial self-propulsion and its relation with cell-length might be elemental to govern the nature of the colony dynamics.

We begin our study by simulating the spatiotemporal dynamics of a collection of rodshaped bacterial cells using an individual-based model as discussed earlier, at a constant packing fraction for varied cell-aspect ratios. Each cell at individual level, posses an identical self-propulsion speed irrespective of their aspect ratio. The packing fraction is taken as *ρ* = 0.65 throughout for the simulation purpose unless otherwise mentioned. Nonetheless, we have also explored the effect of packing fraction on the dynamics of bacterial colony. These motile cells within the colony mutually interact mechanically by elastic repulsive force and move. The images in Figure 2 demonstrates the snapshot of bacterial motion in a simulated colony at a particular time (1836 s) for different cell-aspect ratios (4, 7 and 10). From the snapshot, a qualitative difference in the spatiotemporal organization or colony morphology for different cell aspect ratios is clearly evident. As apparent from Figure 2 and shown in the corresponding videos S1, S2 and S3 [see Supporting Information (SI)], for the case of low aspect ratio (AR = 4) the extent of collective motions are weak, for an intermediate aspect ratio (AR = 7), it shows large scale collective motion dominated by swarming dynamics and finally for larger aspect ratios (AR=10) bacteria start to form lane-like structures and can not rotate properly due to the large cellular length of individuals. In a swarming colony, each corresponding to a specific cell aspect ratio, the flow field fluctuates in both space and time resulting in a distribution of velocities. To quantify the observed features we next calculate the mean speed and vorticity of cells in each colony and statistically analyze the results.

**FIG. 2:**
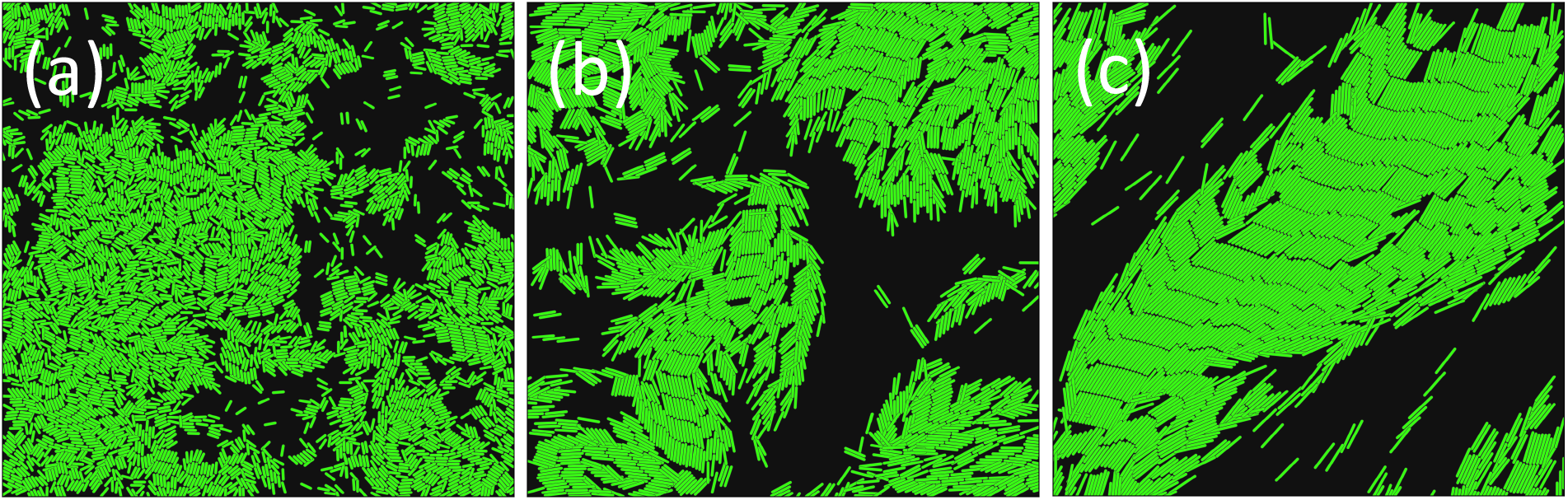
Spatial images of simulated bacteria colony at some time point with a packing fraction of 0.65 for different aspect ratio (AR) of cells: (a) AR = 4, (b) AR = 7 and (c) AR =10. The cells are represented by green spherocylindrical objects and the background is shown in black. All the other parameters are same as given in the Table-I and SI.

### A. Mean speed and vorticity show a non-monotonic variation with respect to the cell-aspect ratio

We determine the mean speed of cells in each colony with a particular aspect ratio averaged over multiple runs. The progression of the resulting mean-speed of the colony as a function of cell aspect ratio, obtained by considering *F_mf_* ∝ *L*, is shown in Figure 3(a). Interestingly, the current proposal of linear dependence of motility force with cell length yields a non-monotonic dependence of the mean speed with cell aspect ratio within the range (4-11) following a maxima at aspect ratio ~7 as shown in Figure 3(a). Since a high aspect ratio larger than 11 is highly unlikely to be achieved by bacterial cells in normal realistic situations, we concentrate mainly on the experimentally observable range [32] of aspect ratios between (4-11).

**FIG. 3:**
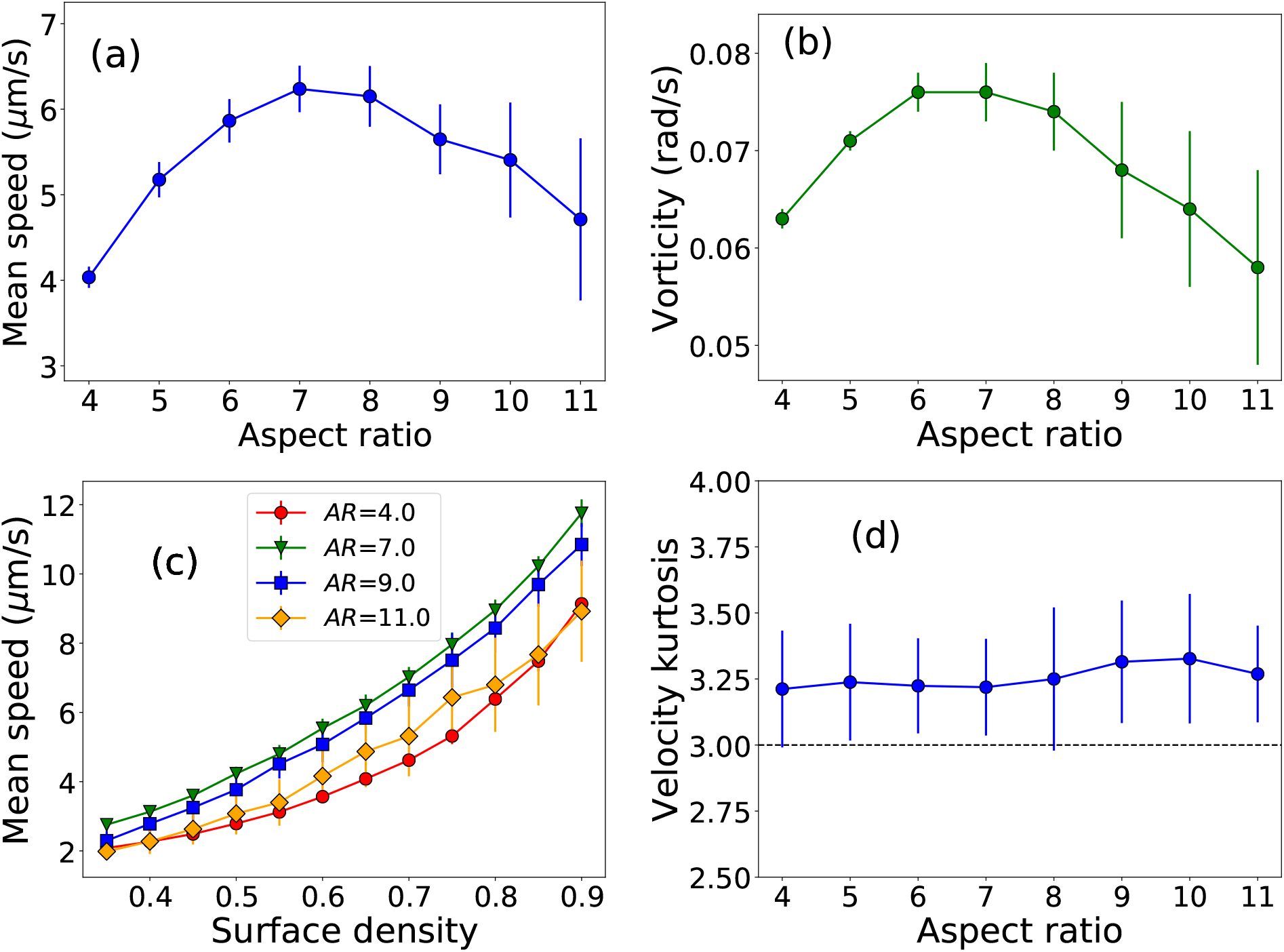
(a) Plot of mean speed as a function of aspect ratio of cell, (b) Plot of vorticity as a function of aspect ratio for a fixed packing fraction *ρ* = 0.65. Both plots show a non-monotonic trend with respect to the cell aspect ratio and show a maximum at AR ~ 7. (c) Plot of mean speed as a function of surface density or packing fraction of cells for different aspect ratios. Mean speed increases with the surface density and the monotonic trend persists for varied aspect ratios. (d) Kurtosis or scaled fourth moment of the velocity as a function of the cell-aspect ratio for a fixed packing fraction *ρ* = 0.65. Error bars here refer to the standard error.

It is apparent from the videos (see supporting information (SI)) that there is some degree of collective swirling motions in these dense dynamical systems of multicellular colonies. In this regard, we compute vorticity, which measures swirls and whirling motions in cellular collectives, at different cell-aspect ratios. We determine 2D-vorticity defined by (*ω* = *∂_x_v_y_* − *∂_y_v_x_*). The mean local flow field 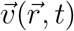 at time *t* and position 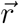 was constructed by binning and averaging individual particle velocities using a spatial resolution. One of the limitations in determining vorticity is spatial resolution. Since we are looking for a comparative observation among vorticity data for different values of cell-size, it is not straightforward because of the different length scales of the cells in each simulated colony. Keeping in mind the largest aspect-ratio value of 11, we chose a bigger spatial grid size of 15 to determine the absolute vorticity. Figure 3(b) demonstrates the plot of absolute vorticity for varied aspect ratios. Vorticity follows the similar traits of non-monotonic trend as observed for mean speed and corroborates fairly with the experimental trend [32].

In order to check how the collective cellular dynamics depends on the surface density of the bacterial cells, we perform a fresh set of simulations by varying the packing fraction of the colony between the range 0.35 — 0.9, thereby maintaining a reasonable high dense organization of cells in an over damped conditions. Figure 3(c) illustrates the plots of mean speed as a function of surface density or packing fraction for different cell aspect ratios. We observe that the mean speed of the bacterial colonies irrespective of the aspect ratio, increases with an increase of surface density, which suggests that as the density of cells increases spatially and locally, they collectively show faster motion [33]. Specifically, we find that for the aspect ratio ~7.0, the mean speed of the colony is largest in comparison to the other aspect ratios. The underlying reason for higher mean speed for larger packing fraction, stems from increasing number of overlaps as number of cells are also higher, which results in high velocities as evident from equations 1 and 2.

The nature of the distribution of velocities in a heterogeneous flow field is known to render further insights into the dynamics of the swarming motion. In what follows, we analyze the fourth moment of velocity called ‘kurtosis’, which characterizes how far the velocity distribution of the whole colony deviates from the normal Gaussian distribution. A value of ‘3’ of kurtosis identifies a perfectly normal distribution. Figure 3(d) demonstrates the scaled fourth-moment (kurtosis), *κ* = *μ*_4_/*σ*^4^, where *μ*_4_ is the centered fourth moment and *σ* refers to the standard deviation, as a function of the cell aspect ratios. We observe that the spatiotemporal dynamics of multicellular bacterial colony follows a Gaussian behavior in their velocity distribution as *κ* ≃ 3 across a wide range of aspect ratios (4-11) for sufficiently high packing fraction (pf=0.65). This trend is qualitatively consistent with previous experimental observation as reported in [33].

### B. Steric repulsive and motility forces together determine the characteristic features of swarming motion

In a dense colony, as the cells move by their self-propulsive forces and eventually interact among themselves mechanically, the effect of steric repulsive force become seemingly relevant. In order to gain insight of how the steric repulsive forces contribute on the swarming dynamics, we now vary the elastic modulus *E* of the cells and carry out the simulations as before. A very high value of *E* ensures perfect hard interaction with almost no overlaps among the cells. However, in a realistic situation, we can only consider a reasonably high value of the elastic repulsive coefficient.

To check the robustness, we performed simulations by varying *E* and repeat each simulation for the entire range of aspect ratios. In Figure 4(a), we plot the scaled(min-max) mean speed as a function of cell-aspect ratio for different values of *E*. As can be seen from Figure 4(a), the variation of elastic repulsive coefficient does not affect the qualitative trend of non-monotonic dependence of mean speed on cell-aspect ratio. To gain a better insight of how the strength of the elastic repulsive force impact on the spatial organization of the overlapping cells in a collection, we varied the power of the overlapping term (*h^s^*) and performed the same set of simulations. It shows that the non-monotonic trend of the mean speed vs aspect ratio curve does not change qualitatively as depicted in Figure 4(b) where mean-speeds are shown following min-max scaling.

**FIG. 4:**
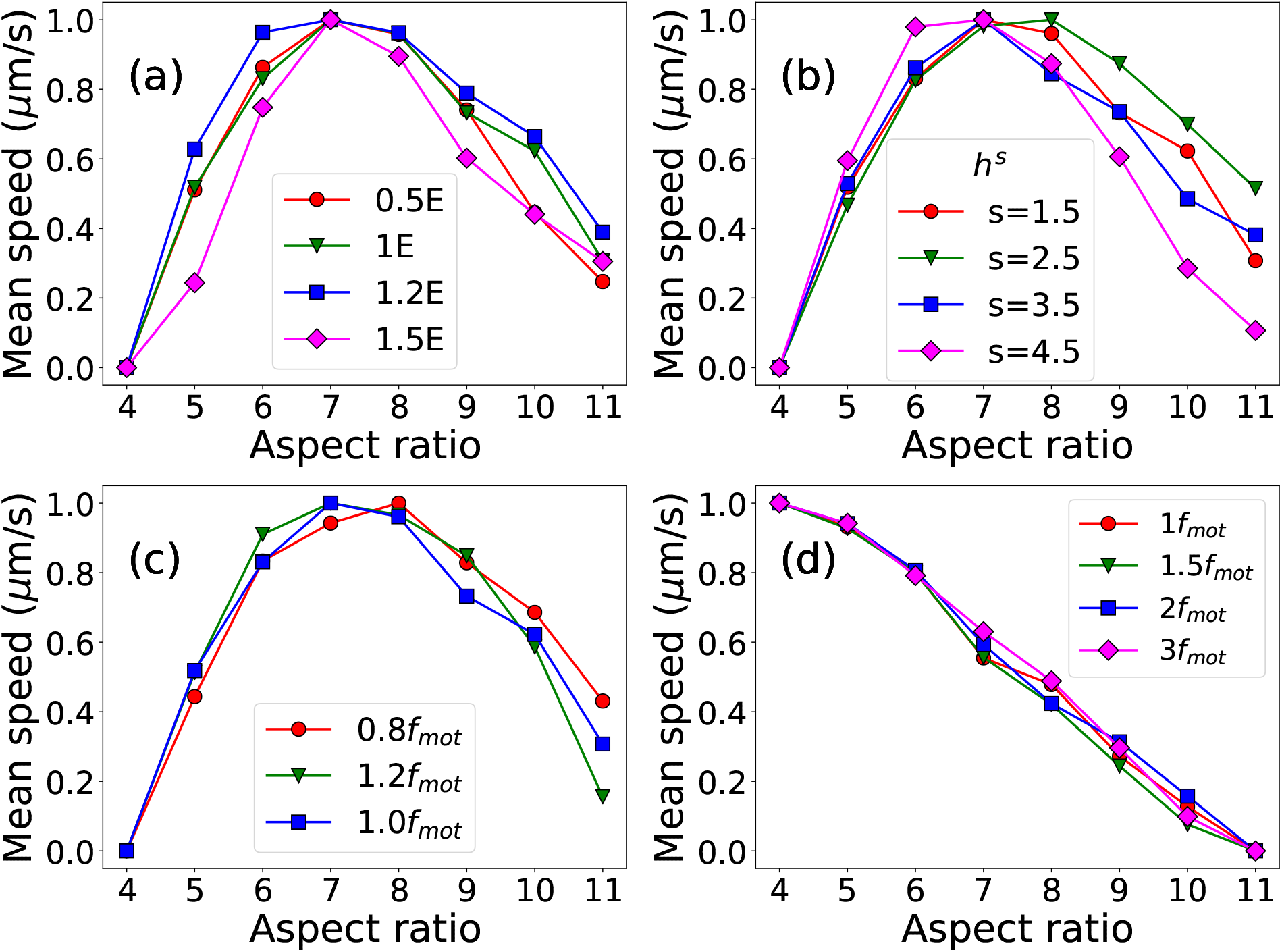
(a)Variation of scaled mean speed as a function of aspect ratio for different elastic repulsive coefficient *E*. (b) Plot of scaled mean speed as a function of aspect ratio for a different power (s) of overlap *h*. (c) Plot of scaled mean speed as a function of aspect ratio for *L*-dependent *F_mf_* with different *f_mot_*. (d) Plot depicts scaled mean speed as a function of aspect ratio for L-independent *F_mf_* with different *f_mot_* values. Non-monotonicity completely vanishes for this case. Here Y-axis of all the figures are scaled between 0 and 1 (min-max scaling).

We next explore the impact of motility force on the collective dynamics of bacteria by varying the motility force and keeping all the other parameter values same as mentioned before. Figure 4(c) demonstrates the variation of the scaled(min-max) mean speed as a function of cell-aspect ratio for several values of *F_mot_* = *f_mot_L*, by varying the value of the constant, *f_mot_*. The result shows a similar qualitative trend of non-monotonic behavior of the mean speed with respect to the cell aspect ratio. These observations infer that the non-monotonic dependence of mean speed on the aspect ratio is quite robust for a range of E and *F_mot_*.

To gain further insights on the roles of *F_rf_* and *F_mf_* on the bacterial collective dynamics, we remove the length dependency from *F_mf_* and take *F_mf_* = *f_mot_*. We varied the *fmot* for different times with respect to the *f_mot_* value given in Table I. Figure 4(d) depicts the variation of the scaled mean speed as a function of cell-aspect ratio for different value of *f_mot_*. We observe a monotonic decay of the mean speed with increase in *f_mot_* which reveals that the non-monotonic trend of mean speed completely vanishes for the aforementioned length-independent *F_mf_*. This observation clearly supports the fact that, motility force and its dependence on cell size plays a crucial role on bacterial motion which leads to the occurrence of the non-monotonic dependence of mean speed on cell-aspect ratio.

### C. Quantitative measures of spatiotemporal dynamics

The overall motion of a cell in a colony mainly depends on two factors: how fast it moves and how long it shows persistent motion in the same direction. In most cases bacteria self-propel following a run and tumble mode of movement using flagella motions. However, prior studies [21, 26], have suggested that mechanical interactions with adjacent cells rather than active reorientation by flagella reversal are the dominant mode of directional changes of cells in a swarm. In order to measure the spatial extent of random motion, we estimate the mean squared displacement (MSD) from the simulated particle trajectories for the different cell aspect ratios. MSD is expressed as MSD(*τ*) = 〈|x(*t* + *τ*) — x(t)^2^〉_*t*_, where x(*t*) refers to the position of cell at time *t* and 〈·〉_*t*_ denotes the average with respect to times *t* for all possible samples. For sufficiently longer times, it is expected that *MSD*(*τ*) ~ *Dτ^δ^* where *D* is the diffusion coefficient and *δ* refers to the exponent of diffusion. From the plot illustrated in Figure. 5(a) we observe that the characteristic exponent δ > 1 for all aspect ratios, implying that the cells are superdiffusive. However, the associated characteristic exponent values differ for varying aspect ratios. We also plot the MSD exponents with respect to the variation of the packing fraction or surface density as depicted in Figure 5(b) which suggests that for both the shorter and longer cells the MSD is lower in comparison to the intermediate cell aspect ratios.

**FIG. 5:**
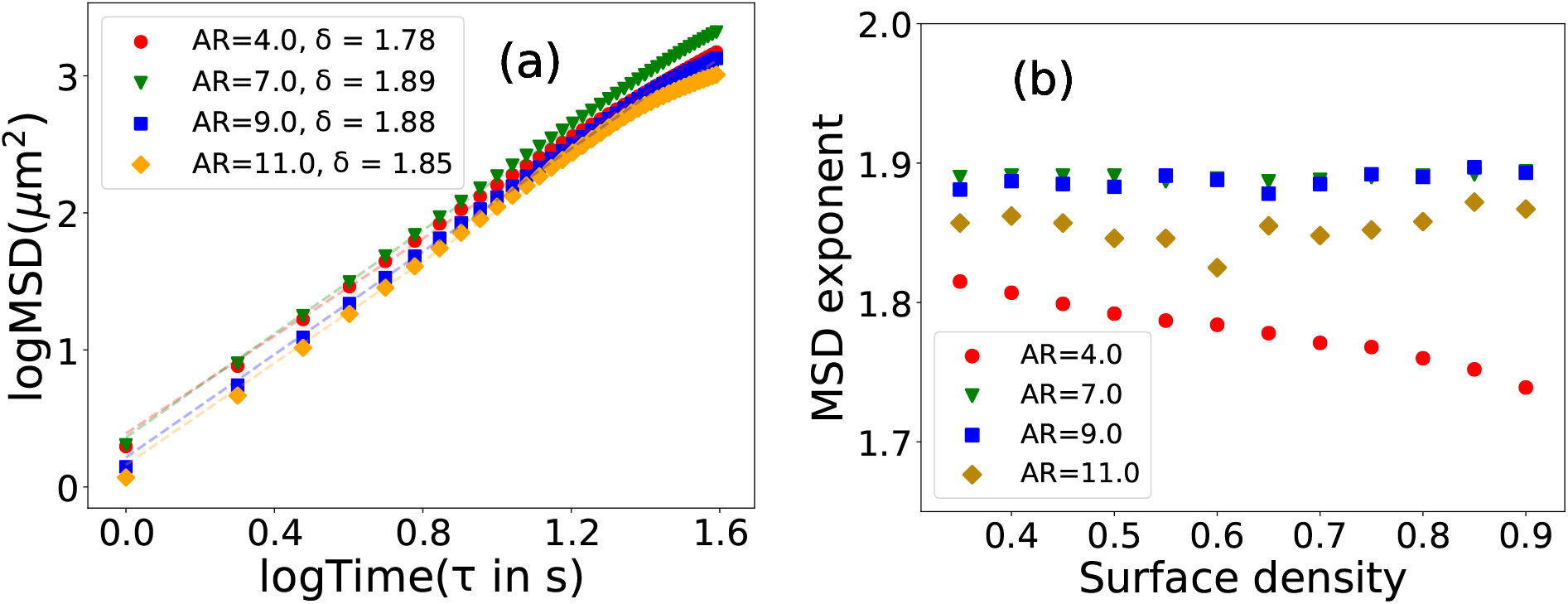
(a) log-log plot of mean squared displacement as a function of time. The fitting curve (MSD(*t*) ~ *D_τ_^δ^*) has been shown as a dashed line. (b) Mean squared displacement (MSD) exponent as a function of surface density for different aspect ratio. Both figure shows that trajectories are supper-diffusive for all aspect ratios and surface densities.

The velocity-velocity time auto-correlation function can provide dynamical insight on swarming motion. The above-mentioned quantity represents the time over which an individual cell’s velocity becomes randomized. To determine time auto-correlation function of velocity, we calculate *C*(*τ*) = 〈*v*(*x,y,t*)*v*(*x,y,t* + *τ*)), where *x, y, t* and *τ* are position, time and lag time respectively and angular brackets denote averaging over time and number of particles. We plot the temporal autocorrelation function of velocity for different cases of cell-aspect ratios in Figure 6(a). It is apparent from the figure, that for a high aspect ratio, temporal auto-correlation of velocity vanishes much slowly. Since, the cells having high aspect ratios in a colony start to form lane-like organized structures, they appear to be strongly correlated.

**FIG. 6:**
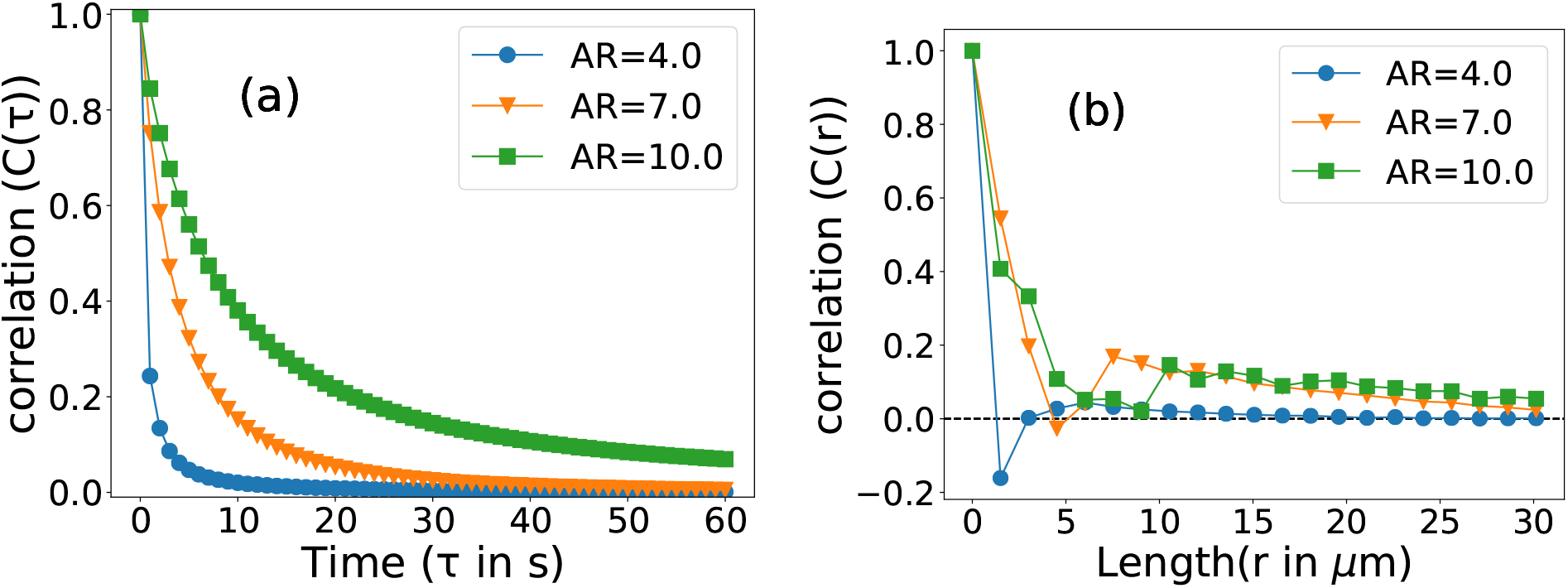
Plot of (a) velocity-velocity temporal auto-correlation function for different values of cell aspect ratios. (b) velocity-velocity spatial auto-correlation function for different values of cell aspect ratios. The other parameters remain the same as given in the Table-I.

To get a closer investigation on how these auto-correlation functions decay over time, we divide the time *τ* into two regimes: one with the initial short time (0-20 s) and the other for the later part of time (21-60 s). These correlation functions for different aspect ratios of cells apparently show a mixed exponential decay over short time and a power law decay over long time. We fit *C*(*τ*) to a double exponential form (for short time) as Ae^−τ/τ_1_^ + (1 — *A*)*e*^−*τ/τ*_2_^, with two time-scales as *τ*_1_, < *τ*_2_ as well as power law form as *at*^−*b*^ for long time. Power law decay implies that there are heavy tailed auto-correlation function for long time due to long range correlation. We also fit the auto-correlation function with a stretched exponential form as *e*^−*αt*^*β*^^ for short time and plot the stretched exponent as a function of cell aspect ratio as shown in Figure-S1 in (SI), which qualitatively resembles with the observation reported in the study [32]. In what follows, our results reveal that two exponential functions are required to fit the time auto-correlation function for all the aspect ratios. Corresponding fitting parameters and plots are given in Table-S2, S3 and S4 and Figure S2, S3 and S4 in the SI.

A bacterial swarm is not only a temporally correlated system but also a spatially coordinated one composed of a large number of cells. Apparently, it shows dynamic patches or groups of cells whose swimming behavior i.e., speed and direction is similar. To quantify these features, we now estimate the correlation between different cells as a function of distance of separation.The spatial correlation function in velocity is expressed as *C*(*r*) = 〈**v**(*x*_1_,*t*).**v**(**x**_2_,*t*)), where **x_1_, x_2_** are two different position and t is time. The angular brackets denote averaging with respect to time and all pairs of points which are separated by *r* = |**x_1_ — x_2_**|. The Figure 6(b) shows the variation of spatial autocorrelation as a function of spatial length. For a small aspect ratio, the alignment interaction is weak among particles and they frequently move in random directions, so their correlation decreases within a short distance. In contrast, for a larger aspect ratio, there is proper alignment among adjacent particles and thus they show finite correlation up to a certain length scale after which it starts to decay. We see that spatial auto-correlation function also shows a long-range correlation for larger aspect ratios.

### D. Asymmetric friction has distinct effects in swarming dynamics

So far we have considered that the parameter friction is same for both parallel and perpendicular directions with respect to the major axis of the cell’s and friction force being proportional to the velocity of the cells, irrespective of the direction of their motion. To understand how and to what extent the friction constants affect the spatiotemporal dynamics in a collection of anisotropic particles, we now consider two different friction constants: ζ_⊥_ (perpendicular direction to cell’s major axis) and ζ_ǁ_ (parallel direction to cell’s major axis). We here define ζ_ǁ_ = *Aζ* and ζ_⊥_ = *ζ/A*, where *ζ* is the previously taken symmetric friction value. We discuss three different cases: (i)*A* > 1 ⇒ ζ_ǁ_ > ζ_⊥_ (rotating motion is easier than translation) (ii) *A* =1 ⇒ ζ_ǁ_ = ζ_⊥_ (symmetric friction) and (iii) *A* < 1 ⇒ ζ_ǁ_ < ζ_⊥_ (translation motion is easier than rotating motion). In case of anisotropy in friction, the friction coefficient can be expressed in the form of a matrix [43] (For details of the derivation see asymmetric friction part in SI). Subsequently the equation of motion becomes modified and can be written as follows

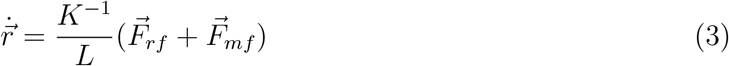

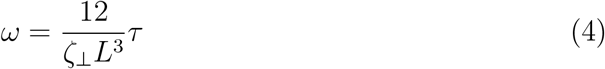

Figure 7(a) and 7(b) demonstrate the variation of mean speed and vorticity as a function of cells aspect ratio respectively, with asymmetric friction. Since vorticity is the measure of local rotation, so for A=1.5 (easy to roll than slide), the cells can rotate more locally compare to A=1 and A=0.90. As angular velocity is proportional to the vorticity, so increasing in angular velocity helps to escalate the mean speed.

**FIG. 7:**
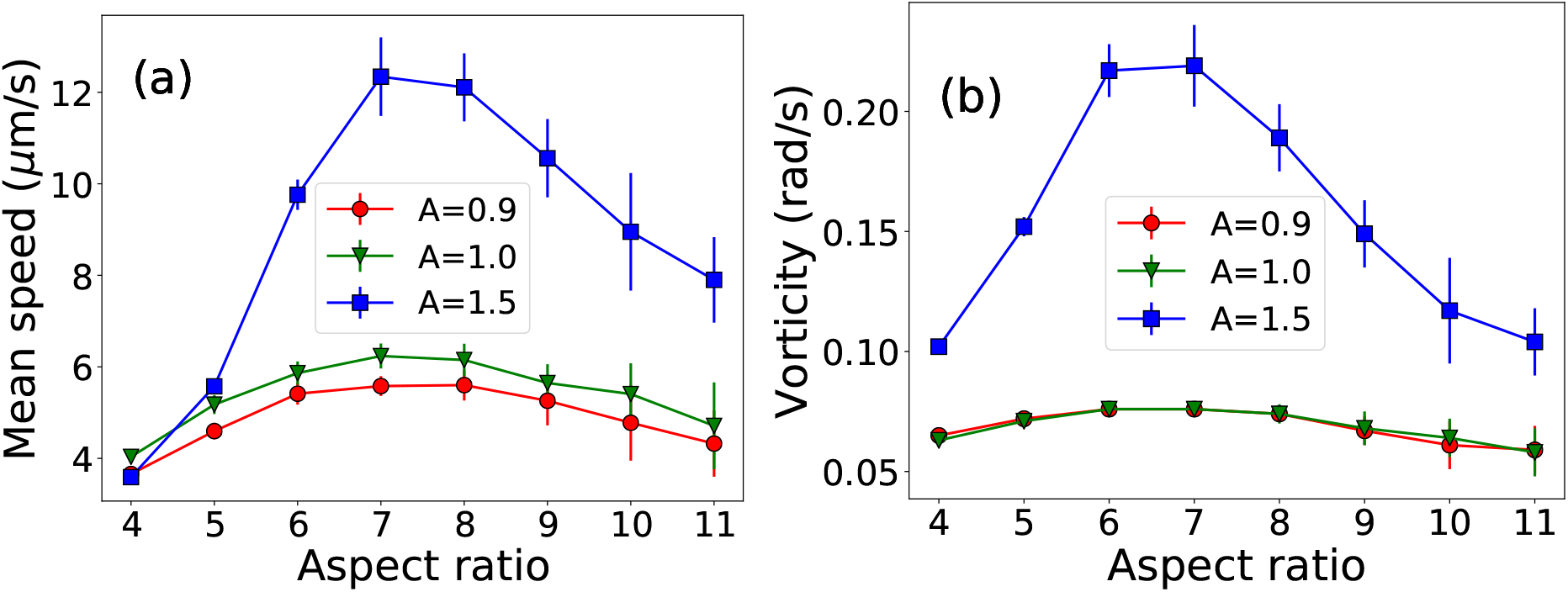
Effect of anisotropic friction coefficients: (a) Mean speed as a function of cell-aspect ratio for asymmetric friction, (b) Vorticity as a function of cell-aspect ratio for asymmetric friction. Both case packing fraction is *ρ* = 0.65. Both velocity and vorticity are high at A=1.5 (easy to roll than slide)

To this end, in order to get further insight on the plausible cause underlying the observed collective dynamics, we estimate the average translational force on each cell experiences which shows a variation with resect to the cell-aspect ratio. Further analysis reveals that for the case of shorter cells, e.g., (AR=4), according to the equations of motion (Eq.(1) and Eq.(2)), the motility force *F_mf_* is small. Moreover, shorter cells lack significant overlap among themselves. These factors cause the net driving force on the shorter cells to be small enough to lower the overall mean speed. On the other hand, for large aspect ratio such as AR=11, the longer cell size and higher motility force render them to align severely to avoid significant overlaps due to collision which results in lane formation as demonstrated in Figure 2(c) and corresponding Movie. When many elongated cells align to form a group of cells, that deflects less easily and therefore moves straight ahead more consistently. This is supported by the lower average force (signified by the large peaks about 0) as shown in Figure-S5 in SI. For an intermediate cell length, the interplay between motility force and the repulsive forces due to overlaps reinforces the total force experienced by the cells in the colony which results in the fastest motions. We found the optimum cell length in our minimal model to be around aspect ratio 7, for which the cells exhibit fastest motion as shown by a maxima in the mean speed and vorticity (Figure 3).

Taken together, these observations lead us to infer that a system of motile asymmetric particles can show variable collective dynamics when the particle size, i.e., the aspect ratio differs. Our observations are quite in line with the existing experimental and theoretical studies on active matter. Interpreting a bacterial swarm monolayer as a purely physical system with cells having size-dependent motility force, we find that the average velocity, vorticity exhibits nonmonotonic dependence on the cell-aspect ratio. To decipher the origin of the aforementioned nonmonotonicity, we have carried out multiple control experiments as discussed earlier, to distinguish between the contributions of both the steric repulsive and the motility forces. We identify that, both short range steric repulsive forces and motility forces jointly decide the fate of the emergent collective dynamics. However, it turns out that the dependence of motility force on cell size is essential for the resulting outcome.

## IV. CONCLUDING REMARKS

Collective coordinated motions of group of biological units is observed across multitudes of scales in living systems. Swarming is one of such collective phenomena, which leads to emerging spatiotemporal behavior in multicellular bacterial colonies. A number of physicochemical processes may contribute to bacterial swarming and these are influenced by cell characteristics as well as environmental conditions [16, 44–47]. In this regard, cell shape and size, is one of such parameters which may lead to variable dynamical scenario in multicellular microbial organization.

Motivated by experimental studies on swarming dynamics of B. *subtilis* [32, 33], in this work, we have explored the connection between the cellular characteristics and physical forces in governing the spatiotemporal dynamics of bacteria collectives with an emphasis to gain a mechanistic insight on the influence of variable cell-aspect ratio in self-propelled rod-shaped bacteria on their swarming motions. We have proposed an agent-based model of motile cells whose self-generated motility force is proportional to the respective cell length. By computer simulations and statistical analysis, we have shown that cell size dependent motility force is a quintessential feature and its interplay with the short-range cell-cell repulsive mechanical force is sufficient to characterize the observed dynamics of flow patterns and statistics of the swarming motion subjected to the variation of cell aspect ratios. Our results predict a nonmonotonic variation of mean speed and vorticity with respect to the cell aspect ratio. For lower aspect ratios mean speed is less, increasing aspect ratio enhances mean speed and it reaches to a maxima around an intermediate aspect ratio where cells show the fastest motions. Further increase in cell aspect ratio leads to a decrease in mean speed. The variation of vorticity also complements the similar trend. These nonmonotonic variations purely stem from the interplay of steric-repulsive forces and a length dependent motility forces. We also explored the role of asymmetric frictions in governing the spatiotemporal dynamics of rod-shaped bacteria which reveals that fastest dynamics will be enhanced if cells can roll than slide in the medium. Our individual based model is able to capture most of the experimentally observed salient properties of swarming dynamics which were not explained by existing theory.

The current investigation has shown that the short range steric repulsive force and cellular length dependent self-propulsive force together is fairly sufficient to understand the dynamics of swarming motion for the bacteria which are only differ by their cell aspect ratio. Long-range hydrodynamic interactions might not be a significant factor to understand the role of cell-aspect ratio on swarming dynamics. Although, the present work is focused on a model bacteria with self-propulsion motion, our study provides a general mechanistic insight of spatiotemporal dynamics of a system of asymmetric rod-shaped particles with the propulsion force dependent on particle size and who are interacting via short-range repulsive mechanical forces. We believe that our study will be relevant to illuminate the features of self-organized spatiotemporal dynamics in similar type of systems where propulsion forces are characteristic function of particle size in the field of soft active matter.

## Supporting information

SI Appendix PDF file in support of the figures: S1, S2, S3, S4, S5 and the tables: S1, S2, S3, S4 as mentioned in the main text.

Movement of bacterial cells for aspect ratio 4 with 130 frames (770-900), separated by 1s. For small aspect ratio it is clear that bacteria are moving

Motion of bacterial cells for aspect ratio 7 with 130 frames (770-900), separated by 1s. Here it is clear that bacterial motion are more alignment

Movement of bacterial cells for aspect ratio 10 with 130 frames (770-900), separated by 1s. For large aspect ratio bacterial cells are forming lanes

## Supporting Information Appendix (SI)

Please see the SI Appendix PDF file in support of the figures: S1, S2, S3, S4, S5 and the tables: S1, S2, S3, S4 as mentioned in the main text. The description of movies S1, S2 and S3 are also included.

## Acknowledgements

We thank N Rana for useful discussions and technical help in coding. All the authors acknowledge TIFR Centre for Interdisciplinary Sciences, India for providing the computing resources. P Ghosh acknowledges the funding from Department of Science and Technology, India in the form of INSPIRE Faculty Award (Grant No: IFA15/CH-201).

