## Supplementary material for "Mechanistic underpinning of nonuniform collective motions in swarming bacteria": SI Appendix PDF file in support of the figures: S1, S2, S3, S4, S5 and the tables: S1, S2, S3, S4 as mentioned in the main text.

### Supplementary Information (Mechanistic underpinning of nonuniform collective motions in swarming bacteria)

#### 1 Simulation and Data Analysis

We have used C language for simulation purposes and calculated all the analyzed quantity using Python language. We have run the program for 32 different initial configurations (ensemble) for total time 2000s with a time step $10^{-3}$s and dumped the data at a separation of $10^3$ time steps i.e 1 s. All the calculated quantities are the ensemble average over 32 initial configurations. For a fixed surface density we have calculated the number of Bacteria using the equation $N = \frac{\rho A}{a}$ where $A = (150 \times 150)\mu m^2$. In Table S1, we have shown the number of Bacteria for a fixed surface density ($\rho = 0.65$). Similarly using this equation, we can calculate the number of bacteria for a fixed aspect ratio with a varying surface density. For all Python scripts we have mainly used Python library like NumPy [1] for numerical calculation, Matplotlib [2] for plotting and SciPy [3] for fitting etc.

#### 2 Fitting Parameter for Autocorrelation

##### 2.1 Stretched exponential

Figure S4 shows stretched exponential ($e^{-\alpha t^{\beta}}$) fitting of temporal auto-correlation function for short time (0 - 20s). Figure S1 depicts the variation of stretching exponent ($\beta$) as a function of cells aspect ratio. All the fitting parameters value is given in the below Table S2

##### 2.2 Double Exponential

Figure S2 shows the fitting of temporal auto-correlation function with double exponential ($Ae^{-\tau/\tau_1} + (1-A)e^{-\tau/\tau_2}$) with $\tau_1 < \tau_2$, for short time (0 - 20s). All the fitting parameters value is given in the below Table S3

##### 2.3 Power Law

Figure S3 shows power law fitting ($at^{-b}$ ) of temporal auto-correlation, for long time (21 - 60s). All the fitting parameters value is given in the below Table S4

### 3 Histogram of force

Here we have calculated the total force per unit length of the cells and plotted their distribution. Figure S5 (a) and (b) shows the distribution of x and y component of total force per unit length respectively, for different aspect ratio. In both cases, the small and large aspect ratio has largely peaked around zero, which implies corresponding this aspect ratio force, as well as the components of velocity, are small, compared to the intermediate aspect ratio.

### 4 Asymmetric Friction

In Eigen basis the form of matrix is $\begin{pmatrix} \zeta_{\|} & 0 \\ 0 & \zeta_{\perp} \end{pmatrix}$,where $K\left|\|\right\rangle = \zeta_{\|}\left|\|\right\rangle$ , $K\left|\perp\right\rangle = \zeta_{\perp}\left|\perp\right\rangle$ and $\left|\|\right\rangle$ , $\left|\perp\right\rangle$ are two Eigen basis.Now any general basis can be written as $\left|1\right\rangle = n_x\left|\|\right\rangle + n_y\left|\perp\right\rangle$ and $\left|2\right\rangle = n_y\left|\|\right\rangle - n_x\left|\perp\right\rangle$, where $n_x = \cos\theta$ and $n_y = \sin\theta$. Now in general basis friction coefficient become

$$K = \begin{bmatrix} \langle 1|K|1\rangle & \langle 1|K|2\rangle \\ \langle 2|K|1\rangle & \langle 2|K|2\rangle \end{bmatrix}$$

which implies

$$K = \begin{bmatrix} \zeta_{\|}n_x^2 + \zeta_{\perp}n_y^2 & \left(\zeta_{\|} - \zeta_{\perp}\right)n_x n_y \\ \left(\zeta_{\|} - \zeta_{\perp}\right)n_x n_y & \zeta_{\perp}n_x^2 + \zeta_{\|}n_y^2 \end{bmatrix}$$

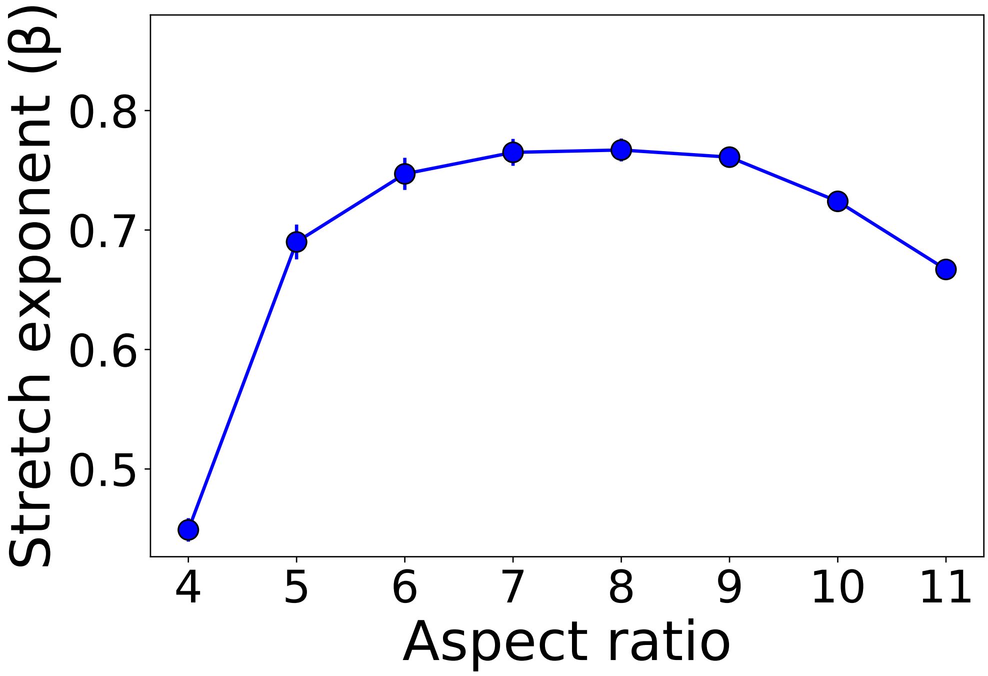


Figure S1: Variation of Stretched exponential ($\beta$)as a function of aspect ratio
.

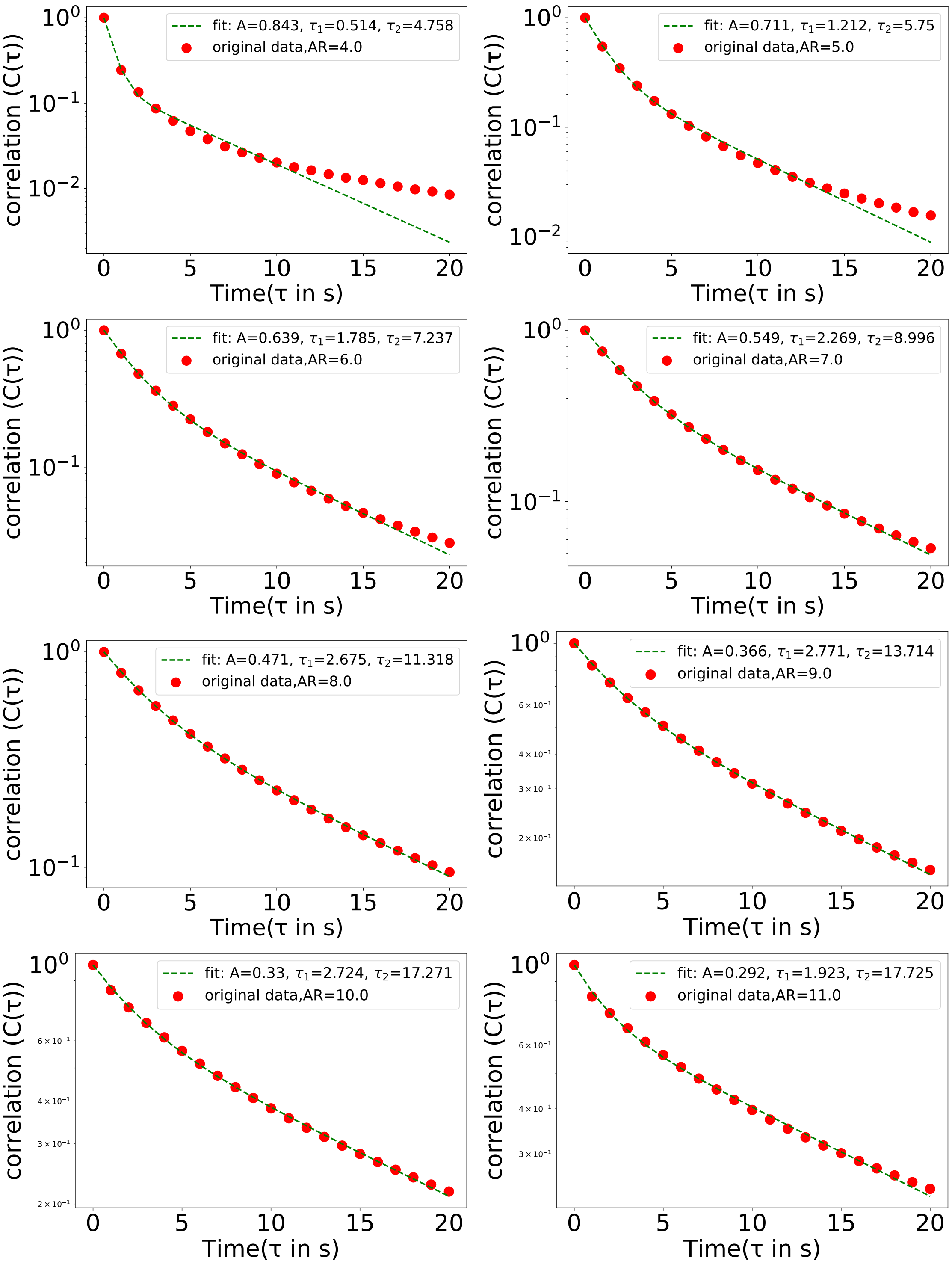


Figure S2: Double exponential fitting for short time

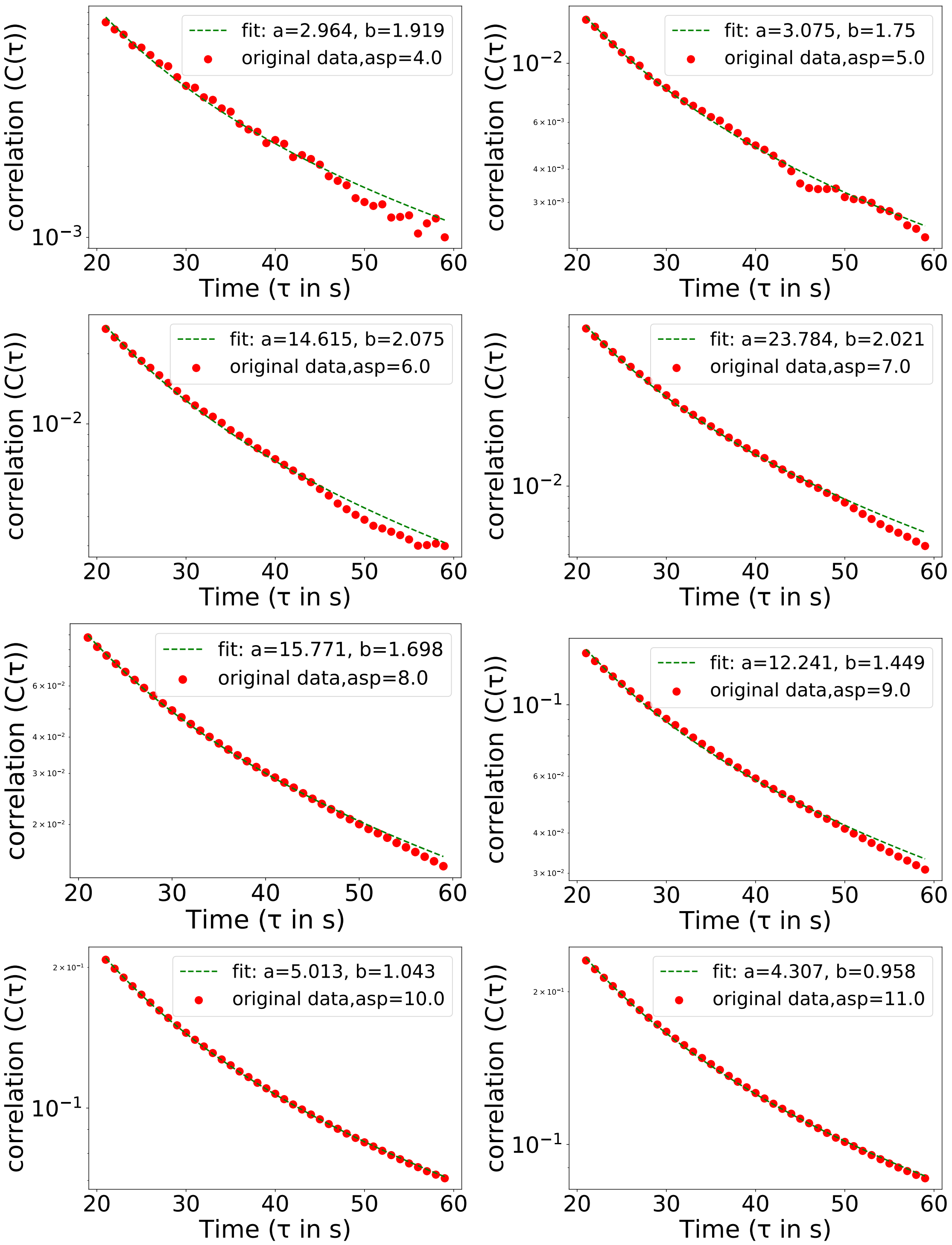


Figure S3: Power law fitting for long time

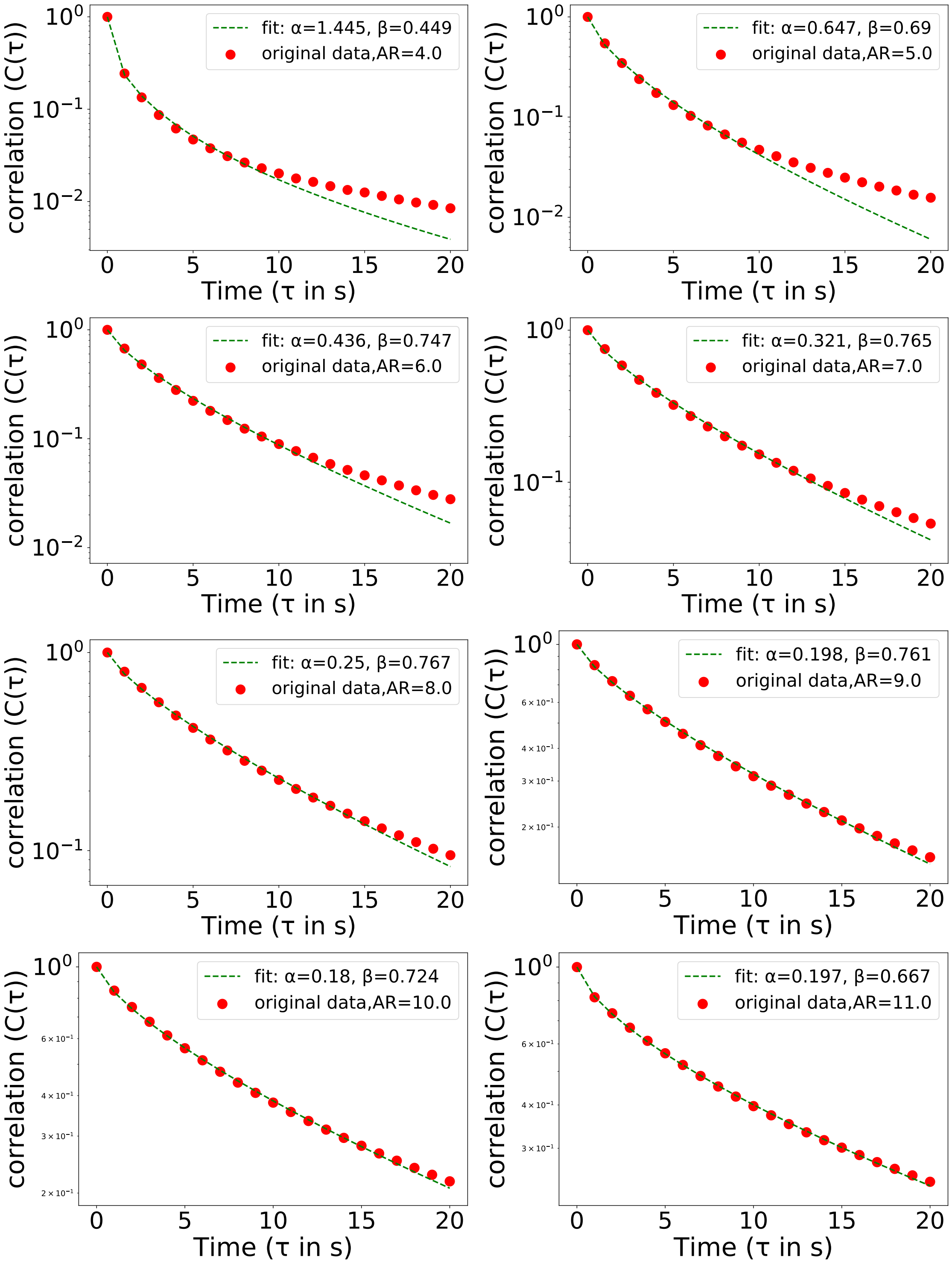


Figure S4: Stretched exponential fitting for short time

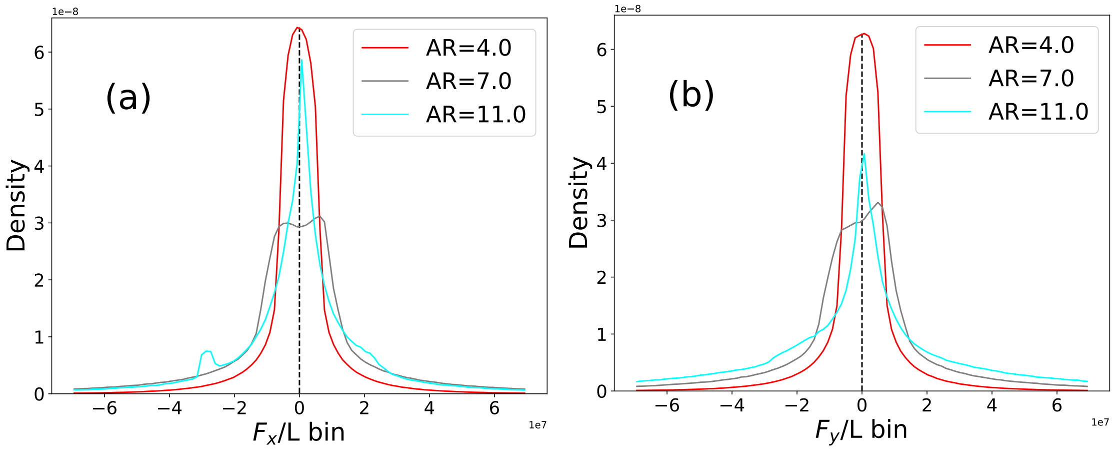


Figure S5: Figure (a) and (b) shows the distribution of x and y component of total force per unit length respectively.

| Aspect ratio | N |
|---|---|
| 4.0 | 3863 |
| 5.0 | 3056 |
| 6.0 | 2527 |
| 7.0 | 2155 |
| 8.0 | 1878 |
| 9.0 | 1664 |
| 10.0 | 1494 |
| 11.0 | 1356 |

Table S1: Number of bacteria for particular surface density

| Aspect ratio | $\alpha$ | $\beta$ |
|---|---|---|
| 4.0 | 1.445 | 0.449 |
| 5.0 | 0.647 | 0.69 |
| 6.0 | 0.436 | 0.747 |
| 7.0 | 0.321 | 0.765 |
| 8.0 | 0.25 | 0.767 |
| 9.0 | 0.198 | 0.761 |
| 10.0 | 0.18 | 0.724 |
| 11.0 | 0.197 | 0.667 |

Table S2: All fitting parameter for stretched exponential fitting.

| Aspect ratio | A | $\tau_1$ | $\tau_2$ |
|---|---|---|---|
| 4.0 | 0.843 | 0.514 | 4.758 |
| 5.0 | 0.711 | 1.212 | 5.75 |
| 6.0 | 0.639 | 1.785 | 7.237 |
| 7.0 | 0.549 | 2.269 | 8.996 |
| 8.0 | 0.471 | 2.675 | 11.318 |
| 9.0 | 0.366 | 2.771 | 13.714 |
| 10.0 | 0.33 | 2.724 | 17.271 |
| 11.0 | 0.292 | 1.923 | 17.725 |

Table S3: All fitting parameter for double exponential fitting.

| Aspect ratio | a | b |
|---|---|---|
| 4.0 | 2.964 | 1.919 |
| 5.0 | 3.075 | 1.75 |
| 6.0 | 14.615 | 2.075 |
| 7.0 | 23.784 | 2.021 |
| 8.0 | 15.771 | 1.698 |
| 9.0 | 12.241 | 1.449 |
| 10.0 | 5.013 | 1.043 |
| 11.0 | 4.307 | 0.958 |

Table S4: All fitting parameter for power law fitting.

### 5 Movie

Movie S1 :- Movement of bacterial cells for aspect ratio 4 with 130 frames (770-900), separated by 1s. For small aspect ratio it is clear that bacteria are moving quite randomly.
Movie S2 :- Motion of bacterial cells for aspect ratio 7 with 130 frames (770-900), separated by 1s. Here it is clear that bacterial motion are more alignment.
Movie S3 :- Movement of bacterial cells for aspect ratio 10 with 130 frames (770-900), separated by 1s. For large aspect ratio bacterial cells are forming lane like structure.
